# Larval seedboxes: a modular and effective tool for scaling coral reef restoration

**DOI:** 10.1101/2025.05.29.656887

**Authors:** Christopher Doropoulos, George Roff, Geoffrey Carlin, Marine Gouezo, Dexter dela Cruz, Aaron Chai, Lauren Hardiman, Lauren Hasson, Damian P Thomson, Peter L Harrison

## Abstract

Natural recovery of degraded coral reefs is constrained by low larval recruitment, limiting restoration at ecologically meaningful scales. While propagule-based approaches have proven effective in plant-dominated systems, scaling larval restoration for sessile invertebrates like corals remains challenging. Traditional coral larval methods rely on net enclosures, restricting impact to small areas (<75 m^2^). We developed and tested a modular, passive larval delivery system – the larval seedbox – to overcome these spatial constraints. Each unit (600 × 500 × 300 mm; 11 kg) enables delayed release of competent larvae near the benthos, enhancing substrate encounter rates over broader areas. At Lizard Island (Great Barrier Reef), five seedboxes delivered ∼14 million larvae across ∼2 ha of degraded reef. Larval release coincided with slack currents to facilitate local retention and subsequent dispersal. Settlement was assessed on 234 tiles placed in concentric arrays around each seedbox. After 48 hours, 85% of tiles had settlers (up to 1,041 per tile), with mean densities 24-times greater than background levels. Enhanced settlement was directly quantified across >470 m^2^, with spatial modelling estimating >3,000 m^2^ via tidally driven dispersal. The larval seedbox enables unrestrained, scalable coral larval seeding and represents a practical advance toward broad-scale reef restoration.

## INTRODUCTION

Coastal marine habitats, such as coral reefs, mangroves, and seagrass beds, are experiencing significant degradation due to anthropogenic pressures including pollution, overfishing, coastal development, and climate change (Duarte et al. 2020). These disturbances disrupt ecological processes and reduce biodiversity, ultimately impairing ecosystem services critical to both marine life and human communities (Barbier et al. 2011). Natural recovery processes, especially the recruitment of juvenile organisms, play a pivotal role in the regeneration of these coastal marine habitats by replenishing populations and reestablishing ecological functions (Caley et al. 1996). However, successful recruitment is highly dependent on habitat quality, connectivity, and the presence of adult broodstock, making degraded environments challenging for recovery without intervention (Orth et al. 2006, Hodgson et al. 2011). Understanding and supporting these natural mechanisms are essential for effective conservation and restoration strategies.

Propagule-based restoration has emerged as an effective technique for large-scale rehabilitation of coastal ecosystems (Vanderklift et al. 2020), particularly for plant-dominated habitats like mangroves and seagrasses. Classic examples include the successful planting of mangrove (primarily *Sonner apetala*) propagules in Southeast Asia and the broadcasting of seagrass seeds (*Zostera marina*) in the eastern USA, where extensive meadows of >1,600 km^2^ and >36 km^2^ have been restored, respectively (Lewis III 2005, Orth et al. 2020). However, the release of propagules has not yielded similar scales of success for invertebrate species, as their complex life histories and dispersal needs (Strathmann 1985, Marshall and Morgan 2011) limit large-scale recruitment through simple propagule release. Instead, invertebrate restoration efforts have traditionally been confined to relatively small-scale interventions such as 0.015 km^2^ for oyster reefs and 0.010 km^2^ for coral reefs (Bayraktarov et al. 2016) due to the use of nets, cages, or artificial substrates required to enhance larval retention and survival (Heyward et al. 2002, Rinkevich 2005, Baggett et al. 2015, Harrison et al. 2021). These constraints highlight the need for strategies that consider life-history traits when applying propagule-based restoration methods.

On coral reefs, the experimental deployment of competent coral larvae using restrained approaches – such as larval enclosures – has successfully enhanced coral settlement rates (Heyward et al. 2002, Suzuki et al. 2012, Edwards et al. 2015, dela Cruz and Harrison 2017, dela Cruz and Harrison 2020, Harrison et al. 2021), and in some cases, has restored breeding populations of mature corals (dela Cruz and Harrison 2017, Harrison et al. 2021). These experiments have demonstrated that directly settling larvae onto reefs can be effective in restoring degraded or larvae-limited systems. However, their application so far been limited to relatively small areas (7-75 m^2^) due to the logistical challenges of deploying tents and nets underwater. To address the issue of scalability, the concept of large-scale coral reef restoration through unrestrained releases of sexually produced coral larvae was modelled as a viable strategy by Doropoulos et al. (2019). This approach aims to mimic natural broadcast spawning reproductive processes (Harrison et al. 1984, Babcock et al. 1986) by releasing competent larvae onto degraded reef areas *en masse* to settle across broad areas of reef through targeted placement of releases (Gouezo et al. 2025b). Despite its theoretical potential, there are currently no published empirical studies demonstrating the success of unrestrained larval releases in reseeding reefs at scales required to be ecologically meaningful (>1ha). The success of such releases is further complicated by species-specific larval competency windows (Randall et al. 2024) interacting with the complex hydrodynamics of coral reef systems, which can disperse larvae unpredictably and reduce the likelihood of successful recruitment (Gouezo et al. 2025a).

Given the complex physical and biological interactions affecting larval propagules (Jones et al. 2009, Randall et al. 2020), slowing down the release and dispersal of competent larvae while maintaining their proximity to the benthos should increase both the likelihood of larvae encountering suitable substrate following their release, and the proportion of larvae competent to settle. This strategy leverages natural hydrodynamic processes to reduce larval loss to advection and improve retention near target restoration areas (Gouezo et al. 2025b), while enhancing local retention through an extended release allows for a wider window of larval competency in multi-species wild spawn collections (Randall et al. 2024). The aims of this study therefore tested whether slowing down the release and dispersal of competent larvae enhanced settlement at a target site could be done using an unrestrained approach to scale the overall spatial footprint of larval settlement to spatial scales greater than 75 m^2^. We utilised a modular system – the larval seedbox – to transfer high densities of competent coral larvae to the benthos, slowly allowing their escape from the seedbox initiated at the beginning of a slack current period to disperse across >2 ha of reef.

## MATERIALS AND METHODS

### Larval seedbox design and deployment

The larval seedboxes are six-sided enclosures (length = 600 mm, width = 500 mm, height = 300 mm), constructed from 6 mm clear polycarbonate sheet. Each box includes eight, 100 mm (4 inch) ports covered with 100 µm plankton mesh to allow water flow into the box while retaining larvae, and an access port on the top to fill the box with larvae (Figure 1a). The seedbox base is also made from 6 mm polycarbonate sheet with 546 CNC drilled holes to allow for slow release of the larvae. Holes were tapered (10.5 mm to 2.5 mm) to reduce the larval swim distance within the narrow section of the hole and potentially guide the larvae toward the substrate (Figure 1b). The seedbox base has guides running along the 500 mm sides. These guides allow a 550 × 500 mm section of 6 mm polycarbonate sheet to slide over the holes to prevent premature larval escape (Figure 1c), with the sheet removed once the seedbox is in position. Four aluminium legs (2 mm thick, 100 mm long) are bolted to the box corners to raise the seedbox 50 mm off the reef surface during deployment (Figure 1d). This elevation also facilitates the removal of the slide out hole cover once the seedbox is in place on the reef.

**Figure 1.**
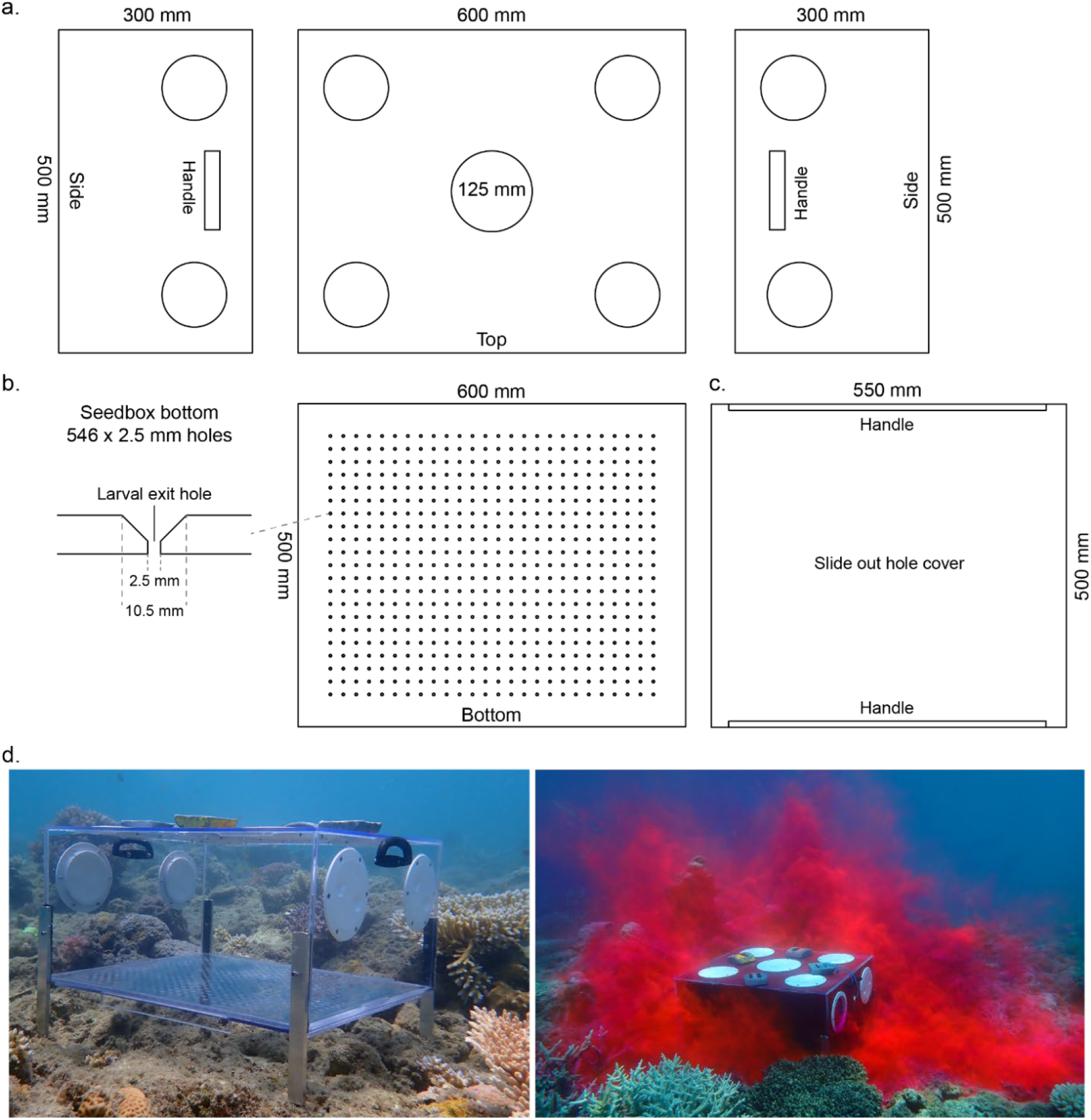
(a) Top and sides of a larval seedbox showing placement of eight, 100 mm holes and a central 125 mm hole on top for inspection ports. (b) Bottom side showing the drill pattern for the 546 holes, each tapering from 10.5 mm to 2.5 mm to encourage larval escape, and (c) the hole cover guides on the base and a 550 × 500 mm cut section of 6 mm polycarbonate sheet that slides over and covers the holes in the base. (d) Larval seedbox (left) prior to larval deployment showing it raised 50 mm from the substrate on aluminium legs and (right) showing the properties of the rhodamine dye coming out from the bottom and dispersing laterally and vertically.

**Figure 2.**
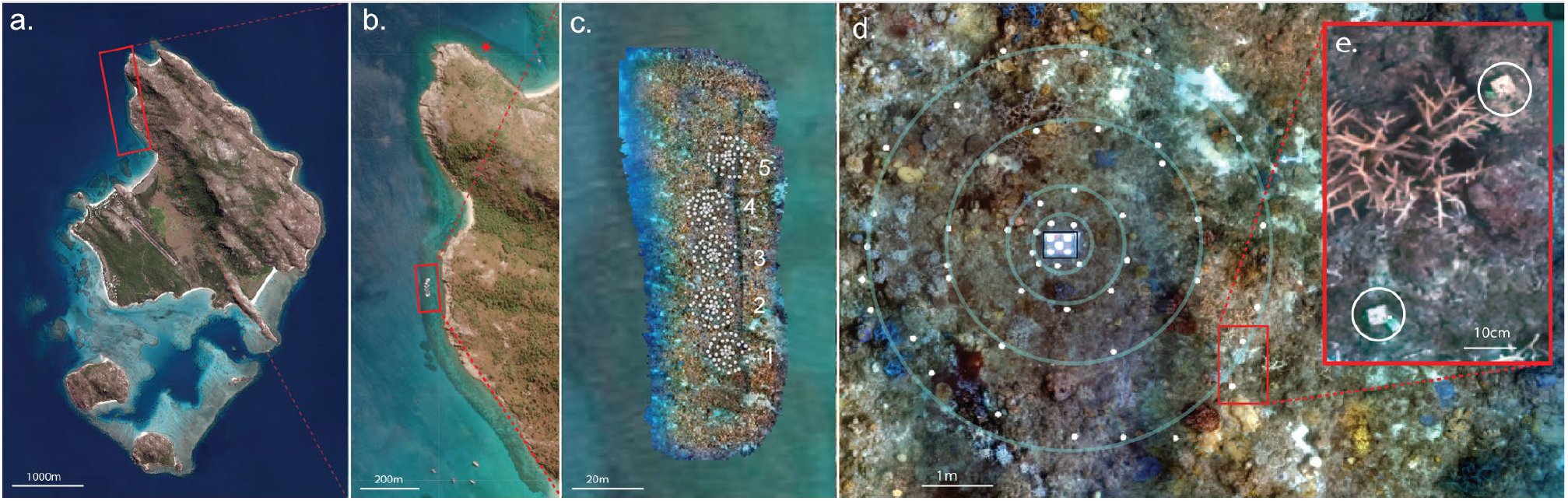
a) Lizard Island, northern Great Barrier Reef, b) study location at Watsons Bay and control location (marked with red star), c) orthomosaic of study location showing the five seedboxes with georeferenced tiles (white circles), d) tile array surrounding each seedbox, e) inset example of georeferenced tiles from orthomosaic.

For deployment, one person lowers the seedbox from the side of a vessel into the water with the central 125 mm inspection port cap removed, then filling it with seawater to about 5 cm from the top. Larval culture was then carefully added into the seedbox through the central port. Any remaining air is allowed to escape before sealing the inspection port. Additional air is assisted out through the 100 mm plankton mesh ports by gentle tapping of the plankton mesh. A single snorkeller swims the seedbox to the deployment site. Once positioned on the reef substrate, up to six dive weights (∼1.4 kg each) are placed on the top of the seedbox to secure it (Figure 1d). Deploying each seedbox takes approximately 5-minutes.

### Larval culturing

Shallow coral reefs surrounding Lizard Island in the northern Great Barrier Reef, experienced extensive coral mortality following the March 2024 coral bleaching event (Australian Institute of Marine Sciences 2024), with an estimated 75% loss of coral biomass overall (Raoult et al *In Press*). However, the reefs located between Turtle Beach and Mermaid Cove in Watsons Bay (Figure 1a), were among the least impacted, retaining approximately 30–40% coral cover. Despite the potential sub-lethal impacts of bleaching on surviving colonies (Briggs et al. 2024), the proportion of gravid *Acropora* colonies in November 2024 (pink/red eggs) was high (Watson’s = 55%, n = 126 colonies; North Point = 67%, n = 162 colonies), indicating good potential for successful gamete collection from wild coral spawn slicks. Therefore, during the predicted spawning window (+2-5 nights after the full moon), collection of wild slicks was focussed along ∼500 m of reef between Turtle Beach and Mermaid Cove, with ∼90 million eggs collected between 2030 and 2300 hours on 20^th^ November 2024 (4 nights after the full moon).

Surface slicks of wild coral spawn were collected using adapted pool scoops (Harrison 2024a), and were distributed among 3 larval culture pools, each measuring 4 × 4 m with a 3 × 3 × 2.3 m 150 µm plankton mesh net (Harrison 2024a). Larvae were cultured in pools for 5.5-days, after which they were observed displaying behaviour indicative of competency i.e., swimming downwards, testing and attaching to the substrate. On the 26^th^ of November, larvae were transferred from one of the culture pools by raising the plankton mesh net and transferring the concentrated larval stock into a 214-litre tub. The larval stock was homogenised by gently stirring the culture to generate dispersive currents within the tub, and samples collected using five 15 ml inverted falcon tubes. Larval density was quantified using stereomicroscopes, estimated at 17 million competent larvae. Approximately 14 million larvae were then distributed equally among the five seedboxes, each box receiving approximately 2.8 million competent larvae.

### Experimental design

Settlement tiles were used to quantify the rate and spatial footprint of enhanced larval settlement. The total area of reef monitored using the tiles was 470 m^2^ (Figure 1c). To detect spatial patterns of settlement surrounding each seedbox, tiles were deployed in a concentric array whereby the inner tiles (n = 6) were located adjacent to the seedbox (<30 cm), and the outer tiles were arranged in three concentric rings at increasing distance from the seedbox (∼1 m, n = 8; ∼2-3 m n = 12; ∼3.5-8 m, n = 24 tiles) (Figure 1d). As the dispersal rate and spatial footprint of larvae were unknown prior to the experiment, the five seedboxes were positioned parallel to the reef slope (Figure 3c) at a similar depth (3.5 m to 5.2 m, depending on tide). Placement location followed the prevailing current direction to optimize detection of spatial patterns within each release, while also allowing for comparisons between seedboxes.

**Figure 3.**
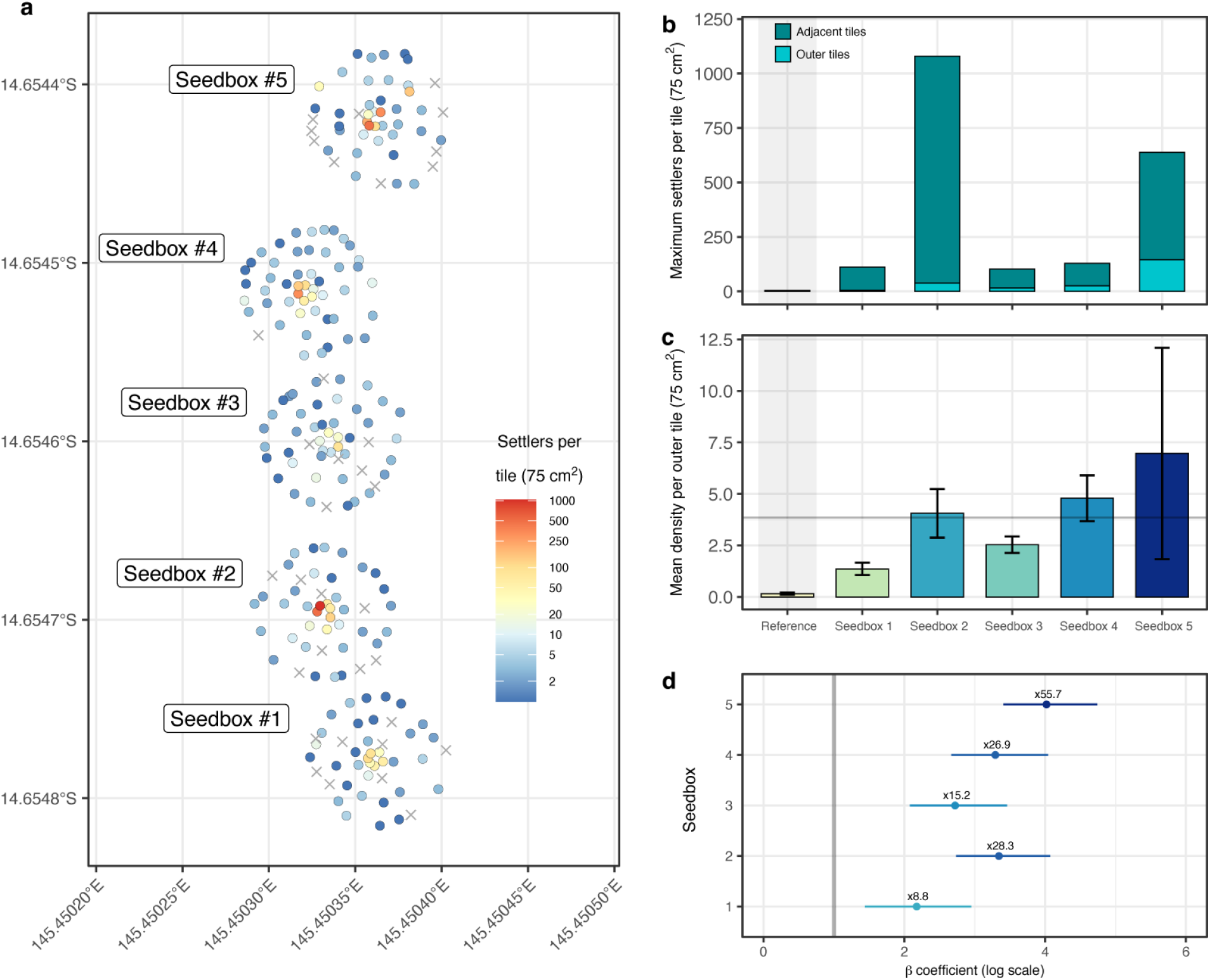
a) Spatial map of tiles within each seedbox array coloured by settler density (where ‘x’ indicates zero settlement); b) maximum settlement density within each seedbox for all tiles (including tiles immediately adjacent to the seedbox) and for outer tiles (>50 cm from each seedbox); c) mean settlement density per tile for the reference site and five seedboxes (horizontal line = mean settlement across all seedboxes, 3.65 larvae tile^-1^); and d) Bayesian posterior median estimates (with 95% credible intervals) from the zero-inflated Poisson model of settlement within seedbox tiles relative to the baseline “natural settlement” at the reference site (with exp(β) or x-fold differences above).

Each tile measured 5 × 5 × 1 cm and was made from travertine, a natural stone formed from calcium carbonate precipitate (Harrison 2024a). Tiles were pre-conditioned for ∼3-months prior to the predicted coral spawning to develop microbial and algal communities that are known to help induce settlement (Heyward and Negri 1999, Webster et al. 2004). Two days prior to the larval releases, tiles were collected, lightly cleaned to remove excess sediment, individually numbered then deployed within the arrays on stainless-steel baseplates with the underside of each tile ∼2 cm from the benthos (following Gouezo et al. (2025a)).

Orthomosaics of the site were generated using a DJI Osmo Action 4 mounted beneath a surface board that was gently towed across the site area, and timestamp correlated to a surface mounted GPS unit (Garmin 65). Resulting photographs were geotagged by writing the GPS into the image exif files, and orthomosaics were generated through Geonadir (www.geonadir.com). Each tile was georeferenced from the image using the ‘sf’ package (Pebesma 2018) in R (R Development Core Team 2024) to allow analysis of spatial patterns in settlement. The total area covered by the settlement tiles spread across 470 m^2^, while the total area covered by the orthomosaic was 2,920 m^2^. Control tiles (n = 63) covering an area of ∼23 m^2^ were deployed at a reference site in similar habitat ∼3 km to the north of the experimental site, allowing for a comparison of settlement rates at our experimental larval release site with a reference site.

On the 26^th^ of November, seedboxes containing ∼2.8 million competent larvae (31 larvae ml^-1^) were deployed between 10:30 and 11:10 am. This deployment occurred just before the predicted slack current, as determined using two tilt current-meters (Lowell TCM-4) and an acoustic Doppler current profiler (ADCP; Nortek Signature 1000) installed at the deployment site. Instruments recorded current data every 5-minutes at depths ranging from 0.3 m to 4 m for several days prior to, during, and after deployment. At 11:30 am, after all seedboxes were deployed, the bottom slide tray and two opposite side windows were removed to allow larvae to disperse onto the reef. Tiles from the experimental and reference sites were collected 48-hours after larval release, and settled corals (i.e., attached and metamorphosed spat) on all tile faces (topside, underside, side) were immediately quantified under dissecting microscopes.

In addition to the *in situ* instrumentation, 60 g of rhodamine dye was added to each larval seedbox three-days later at the identical tidal regime to provide a proxy for the likely dispersal plume of the larval release. The plumes of rhodamine following the opening of the seedboxes were filmed for 300-minutes using a drone (DJI Mini Pro 3) to track the potential dispersal direction and distance of larvae following release. Although the dye trial occurred on a different day to the larval releases, wind conditions remained consistent between the 26 – 30 November. Data from the Lizard Island Automated Marine Weather and Oceanographic Station shows the direction was a consistent south-east to south-south-east but wind strength dropped slightly by ∼25% from a maximum of ∼29 km/h on the 26^th^ to ∼22 km/h on the 29^th^ of November (Appendix S1: Figure S1).

### Analysis of settlement and spatial patterns

Summary statistics of settlement densities within each seedbox were generated using base functions in R (R Development Core Team 2024). Differences in settler counts were modelled across seedboxes using a zero-inflated Poisson (ZIP) regression implemented in the ‘brms’ package (Bürkner 2017). The response variable was the number of settlers per tile, and the predictor was a categorical variable representing treatment levels (seedboxes and “reference” control tiles). The model was fit with four chains, each run for 2000 iterations using default priors. Posterior draws for seedbox-level coefficients (median and 95% credible intervals) were extracted with ‘tidybayes’ (Kay 2020). The posterior median estimates of the log-rate differences (β) between each seedbox and the “reference” settlement site were plotted to show effect size of each seedbox treatment where exp(β) gives the amount higher (e.g. β – 2.94 = 18.9-fold higher than reference).

To assess spatial autocorrelation in settler counts, we performed mark correlation analysis using the ‘spatstat’ package (Baddeley and Turner 2014). Point pattern data were constructed by converting the spatial coordinates of seedbox centroids into planar point patterns (ppp) within a defined observation window derived from the dataset’s bounding box. Settler counts on each tile were assigned as marks to each point. Mark correlation functions (markcorr) were computed with Ripley edge correction over distances from 0 to 6 metres (i.e. maximum diameter of each seedbox tile array) at 20 cm increments separately for the full dataset and a subset of inner tiles using the markcorr function in ‘spatstat’ (Baddeley and Turner 2014). To generate a null expectation for spatial independence of marks, Monte Carlo simulations (n = 999) by permuting settler values across locations while preserving spatial positions. For each simulation, the mark correlation function was recalculated. Simulated values were aggregated, and for each distance the 5th and 99th percentiles were extracted to form an envelope of expected mark correlation under the null hypothesis of random spatial assignment. Observed mark correlation functions were plotted alongside these envelopes to evaluate spatial structuring of settler counts.

To visualize spatial patterns in settler density, a thin plate spline (TPS) model was fit to the outer seedbox tiles (i.e. excluding the tiles in the inner ring adjacent to the seedboxes) through the ‘fields’ package in R (Nychka et al. 2015). Model fitting was controlled by setting the regularization parameter lambda = 100 and using a restricted maximum likelihood (REML), to avoid overfitting and allow for flexible, smooth surface estimation of settler density across the seedbox array.

## RESULTS

Larval release from seedboxes occurred over 24-hours and there were no visible larvae remaining upon retrieval. Aerial drone observations of rhodamine dye releases initiated at the same tidal cycle showed a dense plume persisted over the tile arrays for >4-hours. Approximately one hour after the slack current, a change in current produced a north-northwest flowing rhodamine plume, validated by the *in situ* oceanographic sensors, indicating current-driven larval dispersal.

After 48-hours, larvae had settled on 85.2% of tiles (202 out of 237 total), with densities ranging from 1 to 1041 settlers tile^-1^ (Figure 3a). The highest densities were observed on tiles within 30 cm of the seedboxes (Figure 3b). At distances >50 cm away from seedboxes, the outer array of tiles had variable but substantially lower maximum densities (Figure 3a, b), with a mean density of 3.7 ± 1.5 (± 95% CI) per tile (70 cm^2^). Marked correlation analysis of settler densities on outer tiles showed no evidence of spatial autocorrelation or clustering compared to a null model (Appendix S1: Figure S2), indicating that settlement was randomly distributed across the outer arrays. At a site level, patterns of overall settlement showed an increasing trend from south to north along the reef slope (Figure 3c).

Larval settlement after 48-hours was considerably lower at the reference site compared to the experimental seedbox site, with an average of 0.16 ± 0.10 larvae per tile (± 95% CI), a maximum of 2 larvae settled tile^-1^, and 53 of 63 tiles with no settlers (18% settlement rate). Comparing the seedbox tiles with the reference site indicated that 85% of the tiles at the seedbox site exhibited higher settlement rates than the baseline natural rates. Bayesian posterior median estimates from the zero-inflated Poisson model indicated an 8.8-to 55.7-fold increase in settlement rates within the seedbox arrays, (Figure 3d), averaging a 24.2-fold increase across all outer tile arrays (Appendix S1: Table S1).

## DISCUSSION

The results from this study highlight the potential of the larval seedbox system to significantly improve existing coral restoration approaches by overcoming many of the spatial and logistical constraints inherent in traditional larval restoration techniques (Banaszak et al. 2023, Harrison 2024b). The operational system’s ability to enhance settlement densities on tiles by as much as 56-fold across >470 m^2^ compared to natural background rates demonstrates its potential as a scalable tool for large-scale restoration. These findings align with earlier work conceptualising that harnessing coral spawn slicks presents an opportunity to scale up larval-based restoration (Rinkevich 1995, Doropoulos et al. 2019), provided the challenge of transferring larvae from cultures to the reef at sufficiently large spatial scales can be overcome (Doropoulos 2022, Banaszak et al. 2023, Hughes et al. 2023, Harrison 2024b). The modular design of the seedbox, which allows for slow, targeted larval release that contrasts with more constrained delivery methods (Heyward et al. 2002, Suzuki et al. 2012, Edwards et al. 2015, dela Cruz and Harrison 2017, dela Cruz and Harrison 2020, Harrison et al. 2021), and instead harnesses ambient hydrodynamic processes (Gouezo et al. 2025b) to passively disperse larvae into the benthic boundary layer. This protracted deployment period may also allow larvae to better orient before release and delay exit until reaching a more competent or selective stage, ultimately increasing the likelihood of encountering suitable settlement substrates across broad spatial scales.

The study’s ability to directly observe larval settlement on tiles over a spatial extent of >470m^2^ indicates that this approach could offer a significant advancement over previous methods, which typically restrict the scope of larval seeding to smaller, localized areas (Bayraktarov et al., 2016). While the array of settlement tiles at the larval release site covered 470 m^2^, the orthomosaic covered ∼3,000 m^2^ – which includes the substrate surrounding the tile arrays. Including this available reef substrate and using data from the settlement tiles, modelling indicates that larval settlement densities varied from 250 to 1,250 settlers per m^-2^ (Figure 4a). These densities are similar to the settlement thresholds shown to initiate recovery of juvenile (Gouezo et al. 2021b) and adult corals in experimental work on highly disturbed reefs (dela Cruz and Harrison 2017, dela Cruz and Harrison 2020, Harrison et al. 2021). Over the entire site, larval settlement densities increased from 250 to 1,250 m^-2^ in a south to north direction across the study area, aligning with the rhodamine drift observed from 30-minutes onwards following the release (Figure 4b, c). Moreover, while our tile array and orthomosaic covered areas of ∼470 and 3,000 m^2^, respectively, the rhodamine plume covered a much larger area than the orthomosaic in the initial 30-minutes following release (Figure 4b) and then tracked >1km away in a north-west direction from 30-minutes to 3-hours following release (Figure 4c). ADCP and tilt current meter data shows that the north-north-west dispersal pattern away from the site occurred for 6-12 hours following the release of the larvae. Thus, it is highly likely that coral dispersal and enhanced settlement occurred beyond the immediate study area. Future efforts should aim to create broader arrays of tiles to directly capture the full extent of the settlement originating from the larval seedboxes. This dispersal pattern could enhance the potential for recruitment in areas not directly adjacent to the seedbox, supporting broader restoration goals.

**Figure 4.**
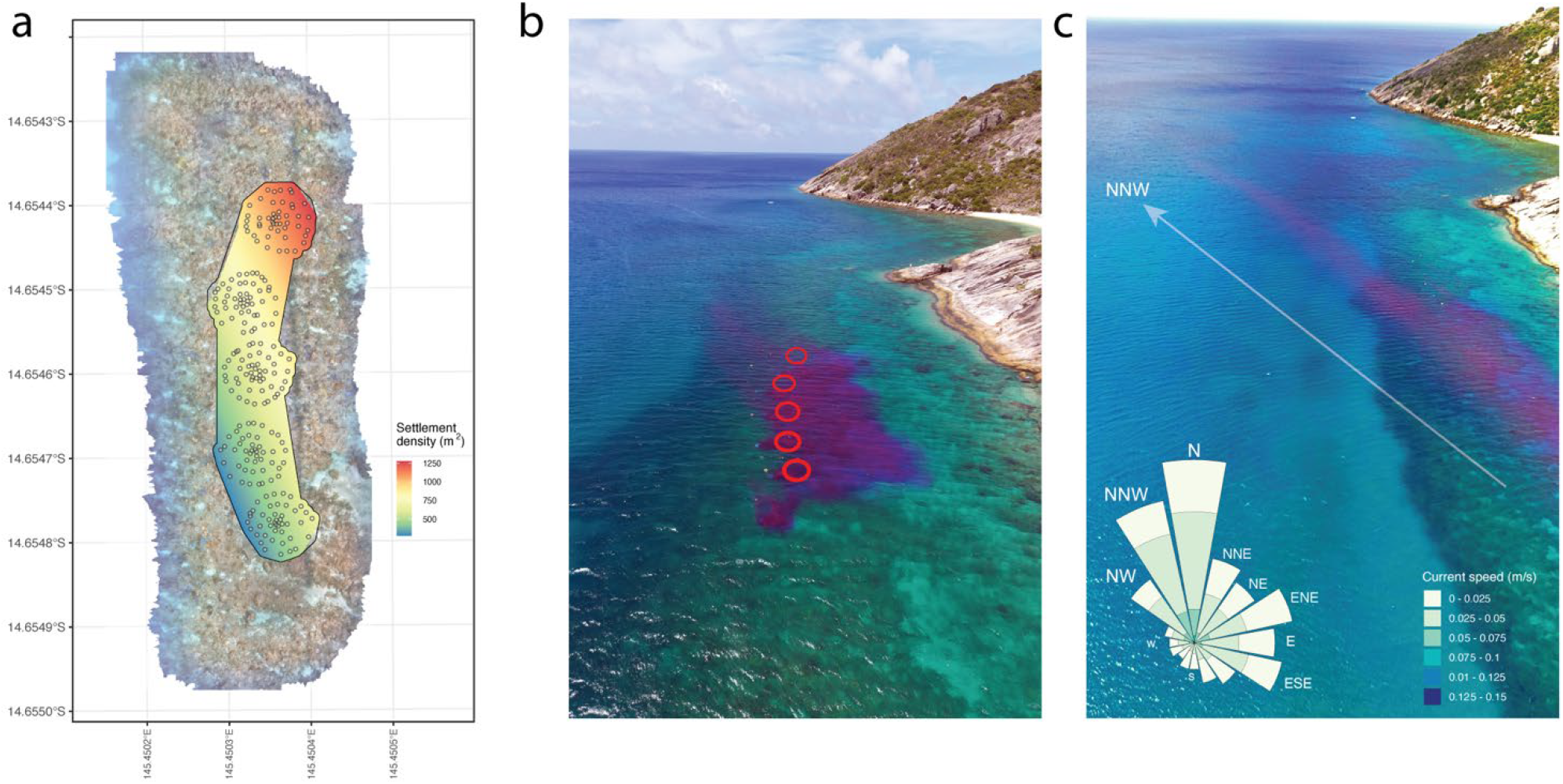
a) Spatial map of seedbox site including orthomosaic, tile arrays, and interpolated thin-plate spline map of settlement density, b), rhodamine cloud 60 minutes after release time c) rhodamine cloud 120 minutes after release time showing NNW dispersal from the seedbox site across adjacent reef habitat, with inset flow data supporting a NNW dispersal of larvae for the 12-hour period following release.

At a highly localised scale, the design of settlement tile arrays provided a precise means of assessing settlement success and any spatial clustering. This approach led to the observation of two clear spatial patterns in settlement density. Firstly, extremely high settlement was observed immediately next to the larval seedboxes. This kind of settlement behaviour aligns with patterns observed from brooding corals that release fully competent larvae – with larvae often settling very close to adults and contributing to highly localised stock-recruitment relationships (Vermeij 2006, Doropoulos et al. 2015). Secondly, the spatial autocorrelation analysis indicated that coral settlement did not form clusters beyond the direct vicinity of the seedbox, supporting the theory that rather than being confined to a small area i.e. 470 m^2^, larvae were dispersed across the broader reef area (>3,000 m^2^). This pattern is likely driven by the movement with tidal-currents and our targeted release time that coincided with the onset of the slack current, which then drifted further away after ∼60-minutes following release. Specific release times utilise natural hydrodynamics for targeted dispersal and larval seeding footprints downstream from release sites (Gouezo et al. 2025b), enhancing the potential for recruitment in areas not directly adjacent to the seedbox to support broader restoration goals of ecosystem scale processes (Gann et al. 2019, Vozzo et al. 2024).

This work has focussed on whether it is possible to observe an increase in coral settlement using an unrestrained release approach at spatial scales larger than previously observed with restrained approaches. It is important to note that settlement on tiles is used to evaluate a short-term signal only, and the study was not designed or intended to investigate any longer-term ecological benefits of population recovery. Further time and monitoring are needed to evaluate long term effectiveness and consideration of location for such studies is extremely important – in particular, locations that are supply limited and have suitable settlement substrate on the reef itself (Gouezo et al. 2021a). For example, some studies have shown successful increase in settlement on tiles and artificial reef structures but no longer term effects on reefs with abundant larval supply and in good condition (Heyward et al. 2002, Edwards et al. 2015), whereas others have found increases in settlement on tiles positively affecting juvenile and adult coral recovery on reef substrates in larval supply limited and degraded reefs (dela Cruz and Harrison 2017, dela Cruz and Harrison 2020, Harrison et al. 2021).

For coral reef management and restoration to be effective, it is essential to consider the specific context and biophysical constraints of the reef habitats targeted for restoration when selecting appropriate tools and techniques (Gouezo et al. 2021a, Quigley et al. 2022). During larval-based coral restoration operations where collection, culture, and delivery all occur *in situ*, the characteristics of the target reef habitat must guide the choice of restoration approach. For example, in well-sheltered shallow reefs protected from wind and waves, anchoring larval pools directly above the reef and allowing cultured larvae to migrate at their own pace through a hose system to the reef is particularly suitable (Heyward et al. 2002, Edwards et al. 2015, Harrison 2024a).

However, this method is limited to sheltered, shallow habitats. In contrast, for reef habitats exposed to wind, waves, and high currents, or where depth limits the safe anchoring of culture pools above the reef for direct transfer of larvae to the reef, the modular seedbox becomes a practical alternative. In these contexts, larvae can be delivered near the substrate using seedboxes – carried out by snorkellers in shallower waters or by SCUBA divers at greater depths – and timed with the onset of slack current conditions to maximize local larval retention following delivery.

The larval seedbox system represents a major advancement in coral restoration by enabling passive larval dispersal during optimal current windows, effectively overcoming key spatial and logistical constraints that have previously limited the scalability of directly seeding larvae to the reef. These results highlight the system’s potential as a practical and scalable tool for large-scale coral restoration, especially in areas where severe reef degradation has diminished natural coral recruitment. The larval seedbox offers a scalable, passive alternative to the initial experimental small-scale net-based coral larval restoration methods, enabling rapid deployment and broad dispersal of competent larvae without the need for complex infrastructure. By overcoming key spatial and logistical constraints – achieving settlement rates up to 56-times higher than natural levels across areas exceeding 3,000 m^2^ – this tool provides a practical, field-ready solution for larger-scale coral reef restoration. This study builds on reviews suggesting that enhancing coral larval survival and settlement through targeted interventions could play a crucial role in restoring coral ecosystems (Randall et al. 2020, Banaszak et al. 2023). Future studies should explore the influence of different larval densities and environmental variables on settlement success (Gouezo et al. 2025a), as well as the long-term resilience of restored populations (Banaszak et al. 2023). Further optimization of the system could increase its effectiveness and applicability in diverse reef ecosystems, offering a valuable resource for coral restoration projects in Indo-Pacific reefs.

## ACKNOWLEDGEMENTS

We acknowledge and thank the Dingaal, Thaanil-Warra and Ngurruumungu Traditional Owners of the Great Barrier Reef for granting free, prior and informed consent (FPIC) to enter and conduct this research on their Sea Country; and thank the Indigenous Partnerships Team at the Australian Institute of Marine Science for their knowledge and time in facilitating FPIC with the relevant Traditional Owner groups. We thank Anthea Donovan and Yvette Millist who managed logistics; North Marine for supporting field operations; Coastal Plastics for engineering the larval seedboxes; and Russ Babcock for reviewing the manuscript prior to submission. This study was conducted under GBRMPA permit no. G20/44511.1.

## AUTHOR CONTRIBUTIONS

Conceptualisation: CD, GR, GC, MG, DdC, PLH; Data collection: GR, GC, MG, DdC, AC, LHar, LHas, DPT; Data analysis: GR; Writing – original draft: CD, GR; Writing – review and editing: GC, MG, DT, PH. Funding acquisition: CD, PLH.

## OPEN RESEARCH STATEMENT

All code, data, and CAD files associated with this manuscript will be freely available at the CSIRO data access portal (https://data.csiro.au/) and https://github.com/marine-ecologist/ upon publication of the manuscript.

## FIGURES

**Figure S1.**
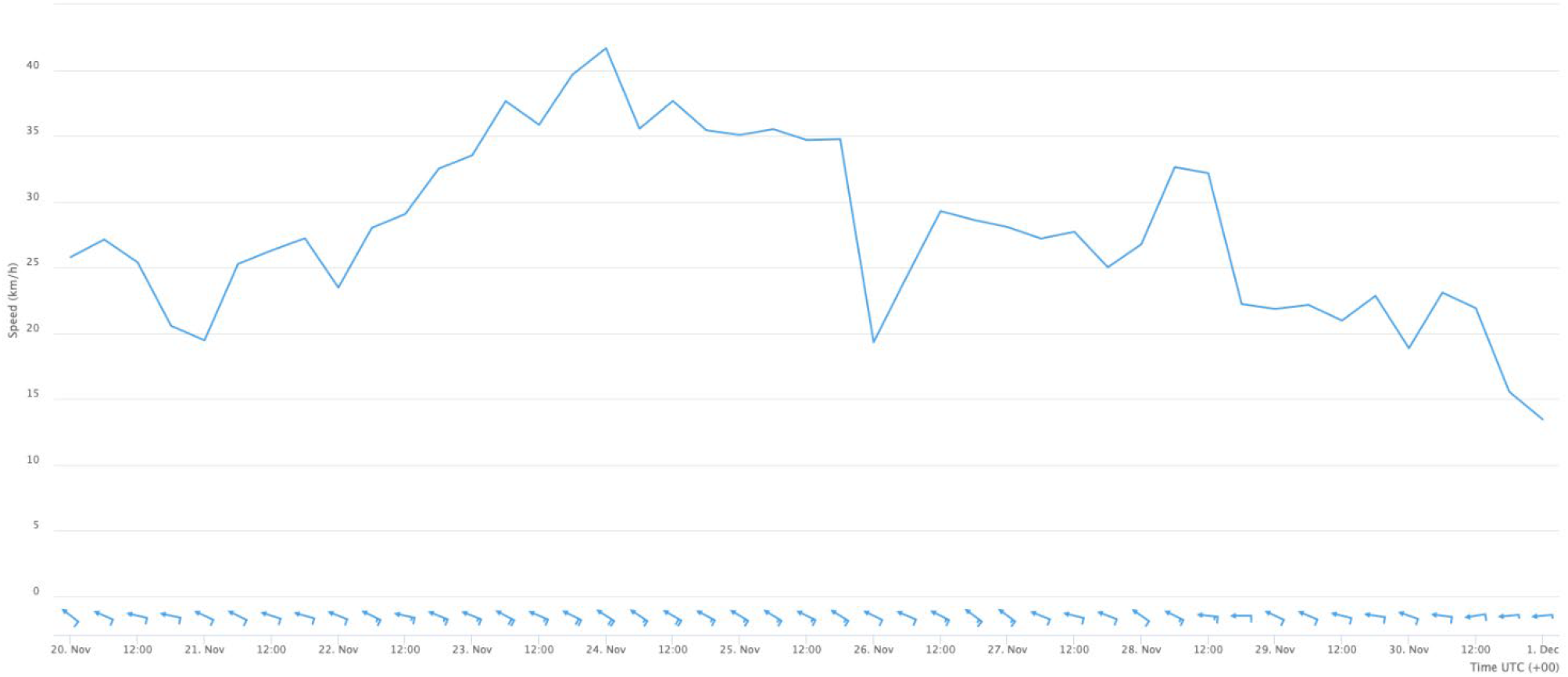
Wind speed and direction data from the Lizard Island Automated Marine Weather and Oceanographic Station sourced from https://apps.aims.gov.au/ts-explorer/?source=3484:Weather%20station&fromDate=2024-11-20T00:00:00&thruDate=2024-12-01T00:00:00.

**Figure S2.**
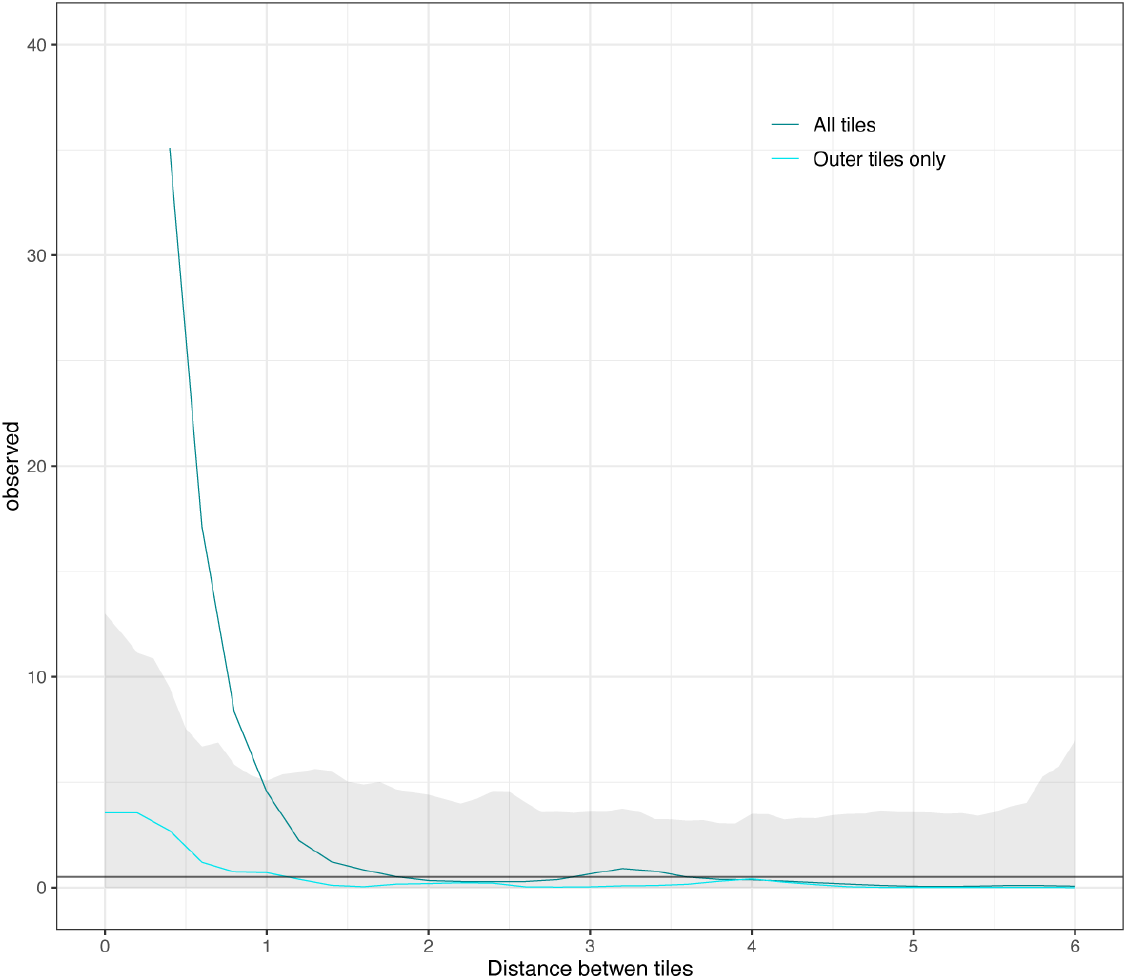
Mark correlation analysis (settler counts per tile as marks to each point) for all tiles and outer tiles, with grey confidence interval representing the 5^th^-95^th^ percent intervals from the of expected mark correlation under the null hypothesis of random spatial assignment.

## SUPPORTING INFORMATION

**Table S1.** Posterior means from Bayesian zero-inflated Poisson model.

| Parameter | Estimate | Error | Lower CI<br>(95%) | Upper CI<br>(95%) | Rhat | Bulk<br>ESS | Tail<br>ESS |
| --- | --- | --- | --- | --- | --- | --- | --- |
| Intercept | -1.71 | 0.33 | -2.4 | -1.1 | 1.0 | 684 | 820 |
| Seedbox1 | 2.18 | 0.38 | 1.44 | 2.95 | 1.0 | 820 | 1026 |
| Seedbox2 | 3.36 | 0.34 | 2.73 | 4.07 | 1.0 | 728 | 825 |
| Seedbox3 | 2.73 | 0.35 | 2.08 | 3.46 | 1.0 | 743 | 828 |
| Seedbox4 | 3.31 | 0.35 | 2.67 | 4.04 | 1.0 | 726 | 960 |
| Seedbox5 | 4.01 | 0.34 | 3.41 | 4.74 | 1.0 | 708 | 868 |
| zi | 0.16 | 0.03 | 0.11 | 0.22 | 1.0 | 2213 | 2169 |

